# *Salmonella* SteD and mammalian SUSD6 use TMEM127 to co-disinhibit the WWP2 E3 ubiquitin ligase

**DOI:** 10.1101/2025.04.23.650309

**Authors:** Samkeliso V. Blundell, Mei Liu, Romina Tocci, Andrea Majstorovic, Solène Tsang, David W. Holden

**Author notes:** Corresponding author –. Indicates current address (Romina Tocci, Andrea Majstorovic, Solène Tsang).

## Abstract

The NEDD4-like E3 ubiquitin ligase, WWP2, is involved in a range of host processes from cell differentiation to T cell immunity. Ligase activity is tightly regulated with WWP2 being held in an autoinhibited state. Binding of a PY motif containing adaptor, an Ndfip, via the WW domains of NEDD4-like E3 ubiquitin ligases leads to their disinhibition. Here, we show that the canonical Ndfip, NDFIP2, requires multiple PY motifs for interaction with and activation of WWP2. In contrast, the single PY-motif containing Ndfips TMEM127 and SUSD6 function as a co-disinhibitory pair. TMEM127 and the *Salmonella* protein SteD also function as a co-disinhibitory pair. However, SteD interacts with a different region of the WWP2, the C2 domain, and this interaction results in disinhibition of WWP2. These findings demonstrate a range of ways that Ndfips can disinhibit WWP2. To our knowledge, these are the first examples of two Ndfips functioning as co-disinhibitory pairs, and of a bacterial effector that disinhibits an E3 ubiquitin ligase.

## Introduction

The NEDD4-like E3 ubiquitin ligases are a family of C-terminal catalytic homologous to E6 COOH (HECT) proteins that catalyse the ubiquitination of themselves and their substrates. They are involved in a wide range of cellular processes, such as the cell cycle, cell differentiation, T cell immunity and viral egress. Their dysregulation can lead to oncogenesis and autoimmunity, highlighting the importance of precise regulation (Maddika *et al*, 2011; Aki *et al*, 2018). They share a common domain architecture comprising an N-terminal C2 domain, two to four tryptophan-tryptophan (WW) domains and a HECT domain. Ligase activity is tightly regulated through auto-inhibition, with disinhibition occurring following binding of an adaptor, binding of calcium or phosphorylation of the E3 ligase (Chen *et al*, 2017; Mund & Pelham, 2009; Wang *et al*, 2010). Other factors also contribute to adaptor-mediated NEDD4-like E3 ubiquitin ligase disinhibition, including membrane tethering, clustering and membrane curvature (Mund & Pelham, 2018; Sakamoto *et al*, 2024). In adaptor-mediated disinhibition, PY motifs (either PPxY, or with one or two prolines in any of the -1, -2 or -3 positions with respect to the tyrosine) within NEDD4 family interacting proteins (Ndfips) interact with ligase WW domains and recruit the ligase to the target protein. Multiple PY-WW domain interactions between either of the canonical adaptors, NDFIP1 or NDFIP2, and the NEDD4-like E3 ubiquitin ligase Itch, leads to release of auto-inhibitory interactions and subsequent disinhibition of Itch (Mund & Pelham, 2009).

The NEDD4-like E3 ubiquitin ligase, WWP2, is co-opted by *Salmonella* effector protein SteD, which is translocated into mammalian cells by the *Salmonella* pathogenicity island-2 (SPI-2) type III injectisome. In antigen-presenting cells, SteD functions via an integral membrane adaptor protein called TMEM127 and WWP2 to deplete mature peptide-loaded Major Histocompatibility Complex class II (mMHCII) molecules, CD97 and B7.2/CD86 from the plasma membrane, thereby inhibiting T cell activation and proliferation (Bayer-Santos *et al*, 2016; Alix *et al*, 2020; Cerny *et al*, 2021). SteD interacts with TMEM127 via amino acids within the SteD transmembrane regions and TMEM127 interacts with WWP2 via the C-terminal PPxY motif of TMEM127: this showed that TMEM127 is an Ndfip (Alix *et al*, 2020). By interacting with both TMEM127 and mMHCII or CD97, SteD induces WWP2-dependent ubiquitination of amino acids in the cytoplasmic regions of mMHCII and CD97, leading to their lysosomal degradation (Alix et al., 2020; Cerny et al., 2021).

Although the interactions between TMEM127, WWP2 and SteD (Bayer-Santos *et al*, 2016; Alix *et al*, 2020) are sufficient to explain how WWP2 comes into close proximity with MHCII, through recruitment by TMEM127 (Alix *et al*, 2020), it is not clear how WWP2 is disinhibited. Interestingly, both TMEM127 and WWP2 were recently shown to deplete surface MHCI in acute myeloid leukaemia (AML) cells. In this case another mammalian protein, SUSD6, which interacts with MHCI and TMEM127, was found to be a component of the TMEM127/WWP2 complex (Chen *et al*, 2023). This raised the possibility that SUSD6 and SteD function with TMEM127 in an analogous fashion. In this study we used mutagenesis, together with cellular binding and disinhibition assays, to shed more light on the mechanism by which WWP2 is disinhibited by a canonical Ndfip (NDFIP2), SUSD6 and SteD.

## Results

### NDFIP2 interacts with the WW2 domain of WWP2

Although WW domains have the greatest affinity for PY motifs in which a tyrosine is preceded by two prolines in the -2 and -3 positions, they can interact with PY motifs that have one or two prolines in any of the -1, -2 or -3 positions (Otte et al., 2003). NDFIP2 is a canonical Ndfip that disinhibits WWP2 (Mund and Pelham, 2009), but the interaction between its three PY motifs and WWP2 has not been reported previously. The first and second PY motifs are of the form PPxY and the third corresponds to LPxY (Fig 1A). First, to determine which WWP2 domains (Fig 1B) are required for the NDFIP2-WWP2 interaction, we constructed 10 truncations of WWP2 extending from either N- or C-termini containing or lacking the C2, WW1, WW2, WW3, WW4 and HECT domains (Fig 1C). NDFIP2 only interacted with WWP2 truncations that contained the WW2 domain (Fig 1C).

**Figure 1.**
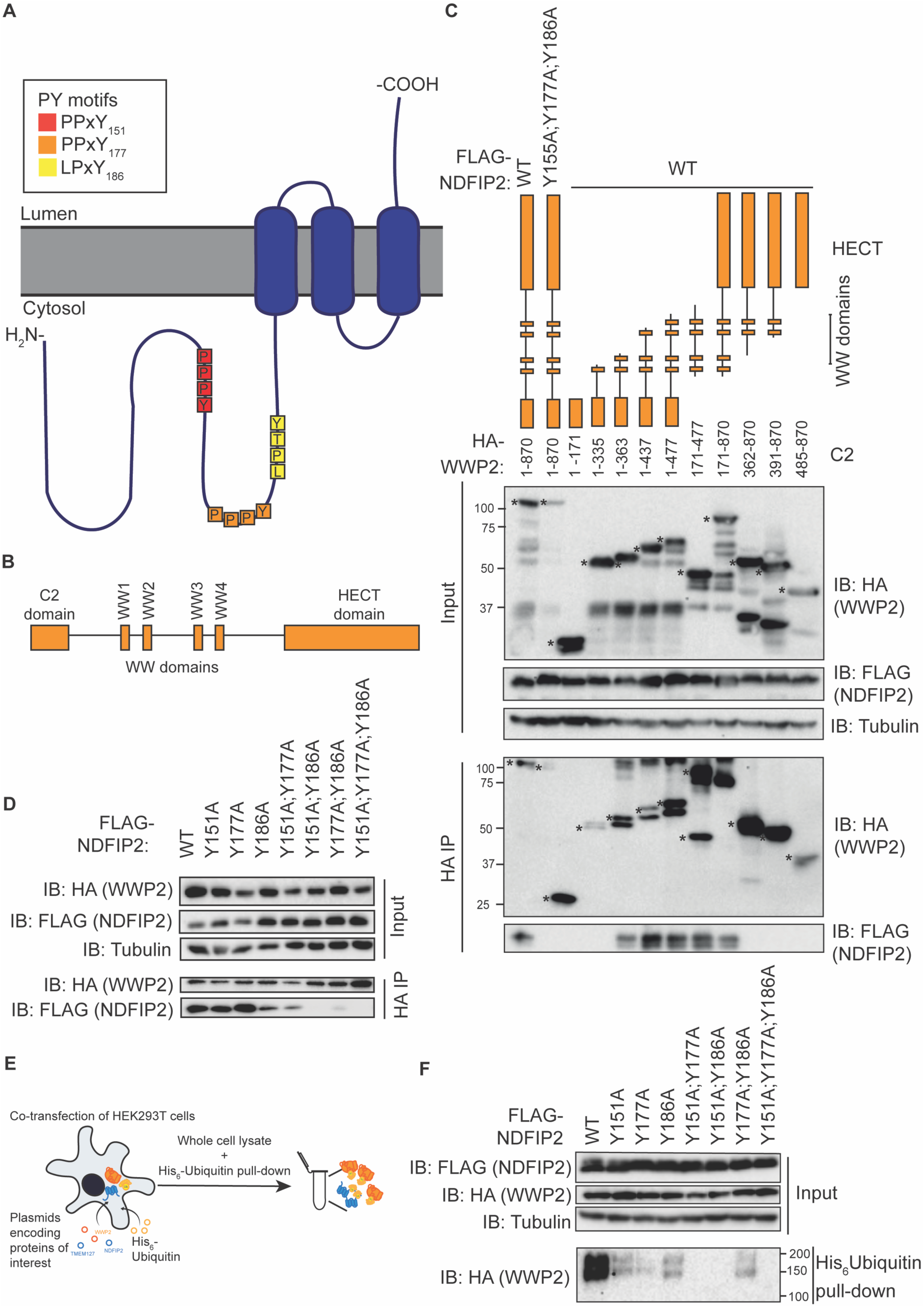
NDFIP2 interacts with the WW2 domain of WWP2 and requires its three PY motifs to interact with and disinhibit WWP2. A. Cartoon of NDFIP2 topology and PY motifs. **B.** Schematic of WWP2 domains. **C & D.** HEK293T cell transfection with FLAG-NDFIP2 (WT or mutant) and HA-WWP2 (full length or truncation as indicated), followed by HA-immunoprecipitation and immunoblot of immunoprecipitate (IP). * Indicates relevant bands. **E.** Schematic of the His_6_-ubiquitin pull down assay used in F. **F.** HEK293T cell transfection with FLAG-NDFIP2 (WT or PY mutant), HA-WWP2 and His_6_-ubiquitin, followed by His pulldown and immunoblot of pulldown. All blots are representative of three independent experiments.

### NDFIP2 requires all three PY motifs to both interact with and disinhibit WWP2

Single, double and triple tyrosine-for-alanine substitution mutants affecting the three NDFIP2 PY motifs were constructed (Fig S1) and assessed for their ability to interact with or disinhibit WWP2. NDFIP2^Y151A^ (PY1-mutated) and NFDIP2^Y177A^ (PY2-mutated) interacted with WWP2 similarly to NDFIP^WT^, whereas the NDFIP2^Y186A^ (PY3-mutated)-WWP2 interaction was weaker and similar to that of the NDFIP2^Y151A;Y177A^-WWP2 interaction (Fig 1D). NDFIP2^Y15A;Y186A^ and NDFIP2^Y151A;Y186A^ had a greatly reduced interaction with WWP2 compared to NDFIP2^WT^ (Fig 1D). Therefore, Y186 of NDFIP2 has the greatest contribution of the three PY motif tyrosines to the interaction with WWP2.

We then assessed the contribution of the PY motifs to disinhibition of WWP2 using a well-established *in cellulo* ubiquitination assay in HEK293T cells (Mund & Pelham, 2009, 2018; Mund *et al*, 2015)(Fig 1E). When WWP2 is disinhibited, it auto-ubiquitinates its HECT domain. Therefore, disinhibition can be assayed by monitoring the ubiquitination of WWP2 (Gong *et al*, 2015). Here, WWP2 and His_6_-tagged ubiquitin were co-expressed with either NDFIP2 (wild-type or mutant as indicated). A His pull-down was then used to assess ubiquitinated proteins. In agreement with previous findings (Mund & Pelham, 2009), NDFIP2^WT^ induced ubiquitination of WWP2 and NDFIP2^Y151A;Y177A;Y186A^ did not (Fig 1F). All single PY motif mutants showed markedly reduced disinhibition of WWP2 compared to wild type. NDFIP2^Y151A;Y177A^ and NDFIP2^Y151;Y186A^ were unable to disinhibit WWP2 (Fig 1F). Evidently, although Y151 and Y177 are not required for binding of NDFIP2 to WWP2, they contribute to the NDFIP2-WWP2 interaction and are important for subsequent WWP2 disinhibition.

### A single PY motif within TMEM127 is required for interaction with WWP2

TMEM127 has three putative PY motifs (Fig 2A): xxPY_220_ of unknown function, PxxY_224_, which together with PPxY_236_ mediates degradation of the tyrosine kinase receptor RET by NEDD4 (Guo *et al*, 2023). PPxY_236_ is also required for TMEM127 to interact with either WWP2 or NEDD4 (Alix *et al*, 2020; Guo *et al*, 2023). We first analysed the contribution of each PY motif and cytoplasmic regions of TMEM127 to SteD-mediated reduction of surface levels of mMHCII during *Salmonella* infection by scanning mutagenesis. Testing of 16 different alanine substitution mutants (Ala1-16) (Fig 2A), each expressed in TMEM127 knockout Mel Juso cells (Fig S2A), revealed only one cytoplasmic region of TMEM127 required for SteD-mediated reduction of surface levels of mMHCII – the Ala16 region (Fig 2B). This region contains the PPxY_236_ motif (Alix *et al*, 2020). Even though the putative PY motifs mutated in Ala13 and Ala14 did not contribute to SteD-mediated reduction of surface levels of mMHCII (Fig 2B), they might participate in ligase interaction. To test this, single, double and triple point mutations of the tyrosine residue within each putative PY motif and the known PPxY motif were generated (Fig S2B) and assessed for their interaction with WWP2. Mutation of Y236 to alanine diminished interaction between TMEM127 and WWP2 with no additive effect upon additional mutation of Y220 and/or Y224 (Fig S2C). Therefore, TMEM127 has a single WWP2-interacting PPxY_236_ motif.

**Figure 2.**
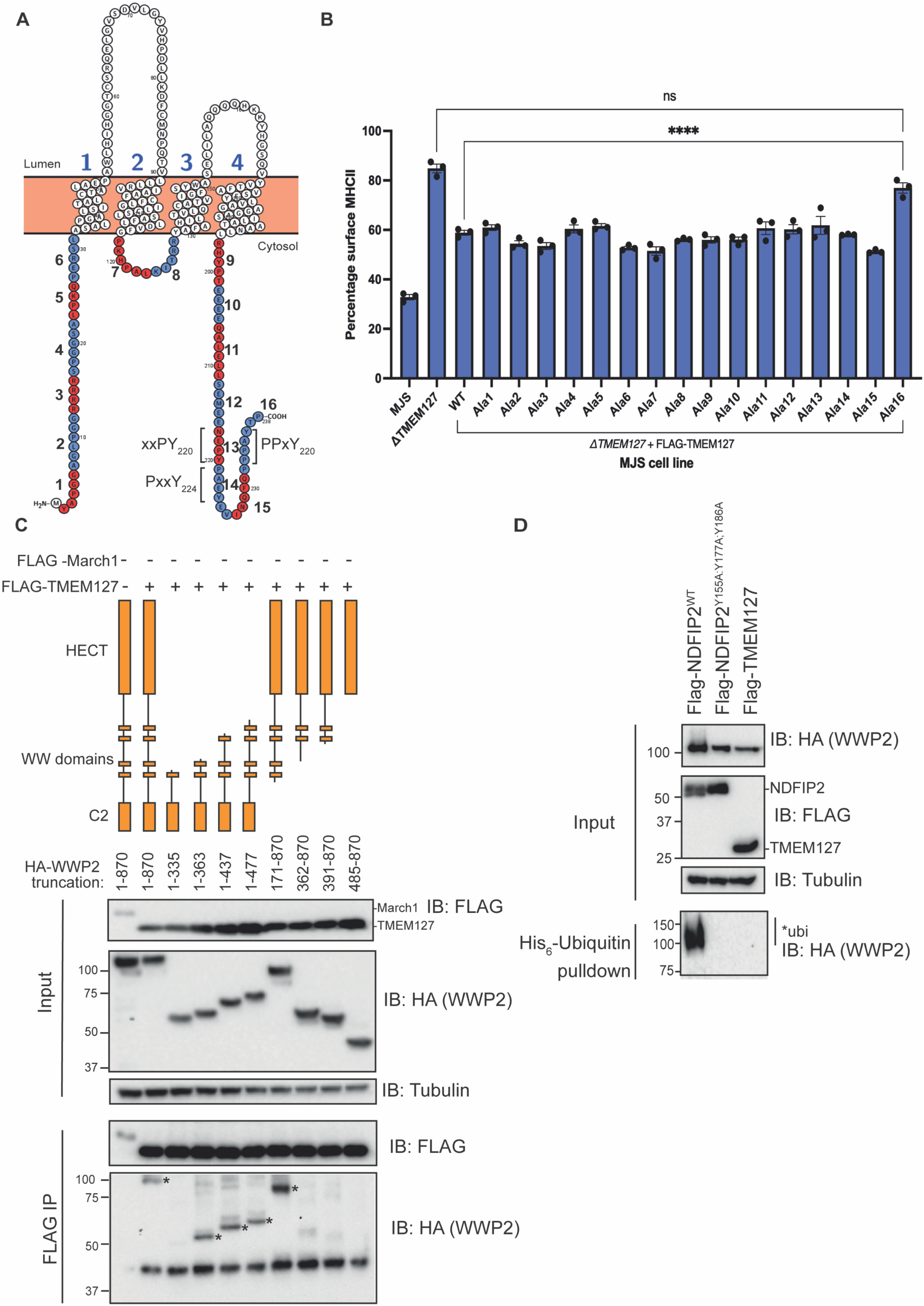
TMEM127 is an atypical non-activating Ndfip of WWP2 and a single PY motif is required for interaction with WWP2. A. Protter image of TMEM127 Alanine mutants highlighted in alternating red and blue (numbered). **B.** mMHCII surface levels of Mel JuSo cells (wild-type or mutant as indicated) infected with *Salmonella*. Cells were analysed by flow cytometry and the data represent geometric mean fluorescence intensity of surface level mMHCII signal of infected cells as a percentage of non-infected cells. Means +/-SEM from three independent experiments. Data were analysed using a one-way ANOVA with a Tukey’s correction for multiple comparisons. N=3. ****p<0.0001 **C**. HEK293T cell transfection with FLAG-March1 or FLAG-TMEM127 and HA-WWP2 (full length or truncation), followed by HA-immunoprecipitation and immunoblot of immunoprecipitate (IP). * Indicates relevant bands. **D.** HEK293T cell transfection with FLAG-NDFIP2^WT^, FLAG-NDFIP2^Y151A;Y177A;Y186A^ or FLAG-TMEM127, HA-WWP2 and His_6_-ubiquitin, followed His-pull down, and analysis of pulldown by immunoblotting. All blots are representative of three independent repeats

### TMEM127 is an atypical non-activating Ndfip of WWP2

Next, we determined the WW domains required for the TMEM127-WWP2 interaction. TMEM127 interacted with WWP2 truncations that contained the WW2 domain (Fig 2C). Therefore, a strong interaction between WWP2 and TMEM127 requires the presence of the WW2 domain of WWP2 and the PPxY236 motif of TMEM127 (Fig 2C), suggesting that these are the principal interacting regions. We then assessed whether TMEM127 could disinhibit WWP2 in the *in cellulo* disinhibition assay. Unlike NDFIP2, TMEM127 failed to induce ubiquitination of WWP2 (Fig 2D), suggesting that it is an atypical Ndfip.

### SUSD6 interacts with WWP2 via a LPxY motif and co-disinhibits WWP2

TMEM127 interacts with an endogenous transmembrane protein, SUSD6, and WWP2 to mediate the reduction of MHCI in tumour cells (Chen *et al*, 2023). SUSD6 has one transmembrane domain and within the cytoplasmic region there is a possible LPxY motif (L_174_P_175_S_176_Y_177_) (Fig 3A). Therefore, we reasoned that SUSD6 might function as an Ndfip of WWP2 and regulate its activity. SUSD6 interacted with WWP2; this was dependent on the presence of Y177 within the LPxY motif and did not require TMEM127 (Fig 3B). Therefore, like TMEM127, SUSD6 is an Ndfip. Although two Ndfips have not previously been shown to function simultaneously on the same NEDD4-like E3 ubiquitin ligase, both SUSD6 and TMEM127 are required for SUSD6-mediated reduction of surface MHCI (Chen *et al*, 2023). Therefore, we hypothesised that SUSD6 and TMEM127 might cooperate by interacting with different WW domains of WWP2. To investigate this, we analysed the domains of WWP2 that SUSD6 interacts with. In addition to interacting with WWP2 truncations containing either the WW2 domain (as for TMEM127 and NDFIP2), SUSD6 also interacted with WW3/WW4 domains (Fig 3C).

**Figure 3.**
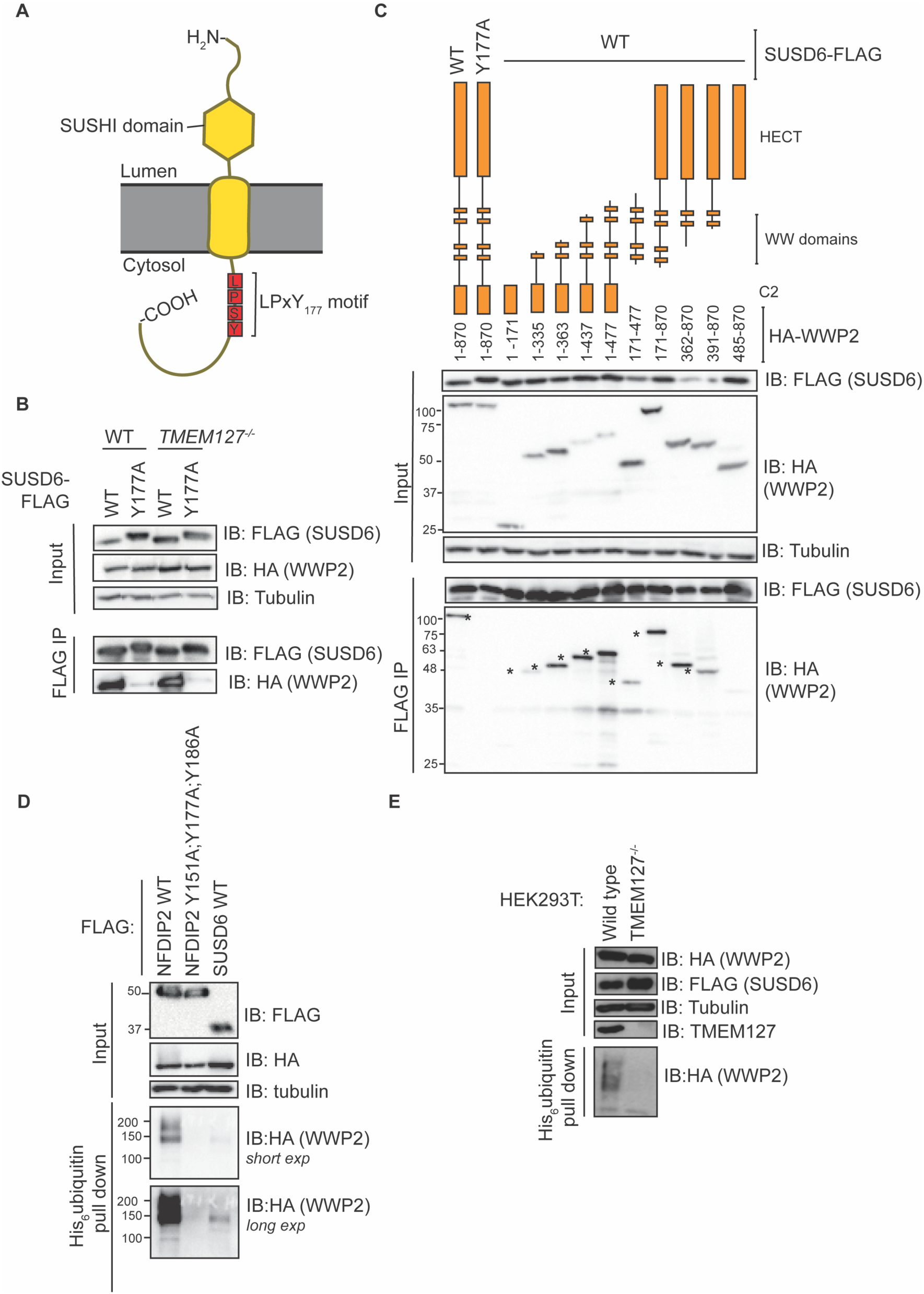
SUSD6 interaction with and disinhibition of WWP2. A. Cartoon of SUSD6 topology and LPxY motif. **B & C**. HEK293T (WT or TMEM127^-/-^) cell transfection with SUSD6-FLAG (WT or Y177A) and HA-WWP2 (full length or truncation) followed by FLAG-immunoprecipitation and immunoblot of immunoprecipitate (IP). *Indicates relevant bands **D & E.** HEK293T (WT or TMEM127^-/-^) cell transfection with FLAG-NDFIP2 (WT or PY mutant) or SUS6-FLAG, HA-WWP2 and His_6_-ubiquitin followed by His pull down and analysis of pulldown by immunoblotting. All blots are representative of three independent experiments.

We then determined if SUSD6 could disinhibit WWP2. In the presence of TMEM127, SUSD6 induced the ubiquitination of WWP2 (Fig 3D). This was less than the amount of ubiquitination induced by NDFIP2^WT^ but greater than the amount induced by NDFIP2^Y151A;Y177A;Y186A^ (Fig 3D). In the absence of TMEM127, SUSD6 failed to induce ubiquitination of WWP2 (Fig 3D). Together, these results show that SUSD6 and TMEM127 co-disinhibit WWP2.

### SteD and TMEM127 co-disinhibit WWP2

Having established the regions required by three endogenous transmembrane Ndfips (NDFIP2, TMEM127 and SUSD6) to interact with and disinhibit WWP2, we investigated how the bacterial effector and transmembrane protein SteD (Fig 4A) contributes to WWP2 function. First, we determined whether SteD could interact with WWP2 and if this interaction required TMEM127. Immunoprecipitation of GFP-SteD from transfected HEK293 cells showed that WWP2 interacted with GFP-SteD; this was reduced but not eliminated in the absence of TMEM127 (Fig 4B).

**Figure 4.**
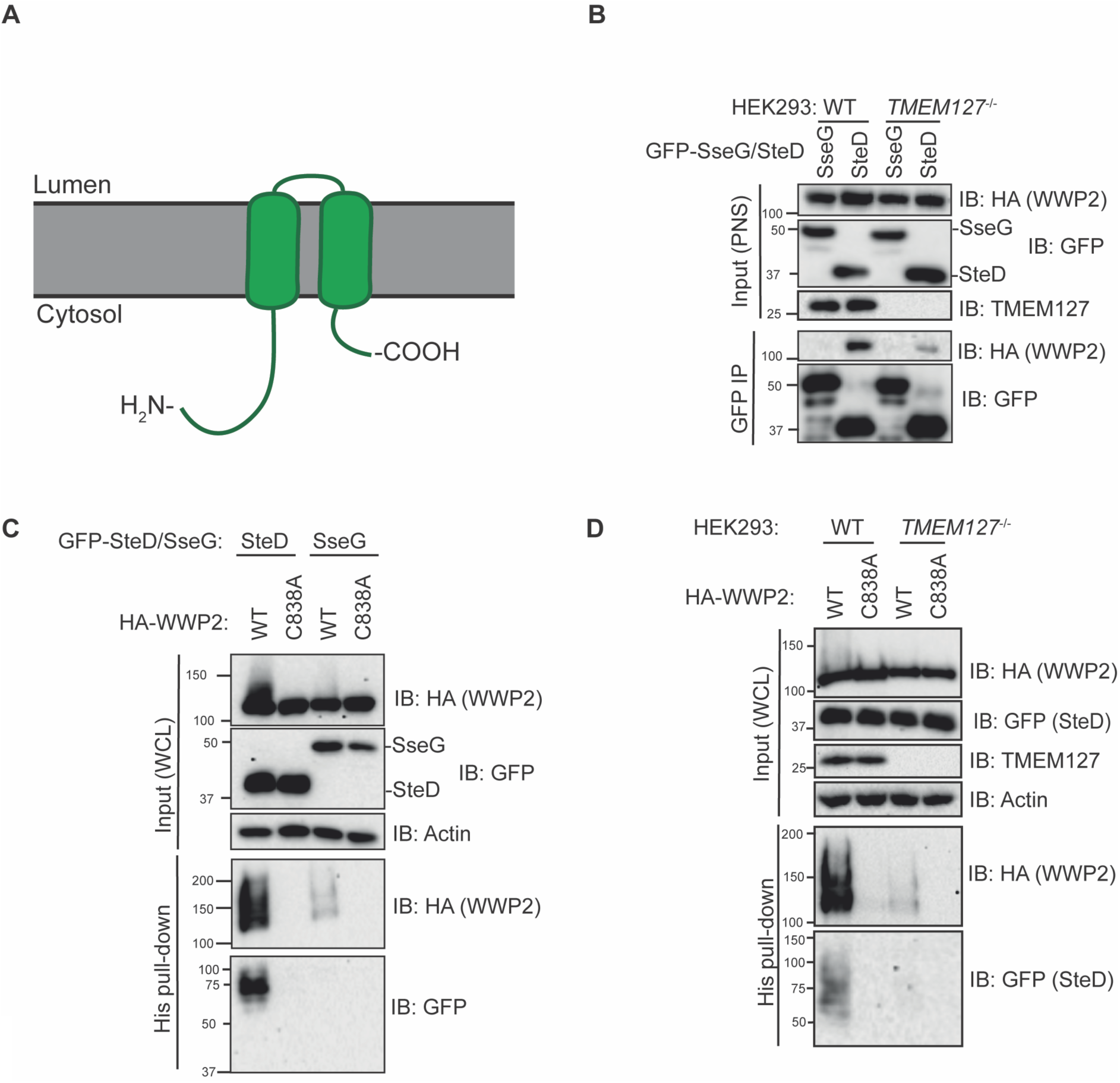
SteD is an NDFIP and TMEM127 and SteD are co-disinhibitors of WWP2. A. Cartoon of SteD topology. **B.** HEK239T cell (WT or TMEM127^-/-^) transfection with GFP-SseG or GFP-SteD and HA-WWP2, followed by GFP-immunoprecipitation and analysis of immunoprecipitate (IP) by immunoblotting. **B & C.** HEK293T cells (WT or TMEM127^-/-^) transfection with GFP-SseG or GFP-SteD, HA-WWP2 (WT or C383A), followed by His pulldown and analysis of pulldown by immunoblotting. Post-nuclear supernatant (PNS); whole cell lysate (WCL). All blots are representative of three independent experiments.

Given that SteD functions with TMEM127 to reduce surface levels of MHCII in a way that is analogous to SUSD6/TMEM127/WWP2-mediated reduction of surface levels of MHCI (Chen *et al*, 2023; Alix *et al*, 2020), we hypothesised that SteD might disinhibit WWP2. Using the His_6_-ubiquitin pull-down assay, we found that SteD induced strong ubiquitination of WWP2, in comparison to another SPI-2 integral membrane effector of unrelated function, SseG (Fig 4C). The very weak disinhibition of WWP2 in cells expressing SseG (Fig 4B) is presumably caused by other endogenous Ndfips in HEK293T cells. SteD also undergoes ubiquitination, and this is important for its function in reducing mMHCII surface levels (Alix *et al*, 2020).

Therefore, we used a catalytic point mutant of WWP2 (WWP2^C838A^) to determine whether SteD-induced ubiquitination of WWP2 was auto-ubiquitination, and if ubiquitination of SteD was caused by catalytic activity of WWP2 (Fig 4C). SteD-induced WWP2 ubiquitination required the catalytic activity of WWP2 and the ubiquitination of SteD itself also required WWP2 catalytic activity (Fig 4C). These results reveal that SteD disinhibits WWP2 and that SteD is ubiquitinated by WWP2.

SteD requires TMEM127 for SteD-meditated ubiquitination of mMHCII (Alix *et al*, 2020). Therefore, we therefore investigated the level of auto-ubiquitination of WWP2 induced by SteD in HEK293T cells lacking TMEM127. SteD-induced WWP2 ubiquitination was greatly diminished in TMEM127 knockouts when compared to WT cells (Fig 4D). Together, these results show that, as for SUSD6 and TMEM127, SteD and TMEM127 work together as co-disinhibitors of WWP2.

### SteD interacts with TMEM127 and WWP2 and requires its N and C-terminal cytosolic domains to disinhibit WWP2

In addition to lacking a PPxY motif, SteD lacks proline-rich regions and any putative PY motifs that could interact with WW domains and contribute to its ability to disinhibit WWP2. Therefore, to determine which region/s of SteD contribute/s to the interaction with and disinhibition of WWP2, truncations of SteD were used (Fig 5A and S3)(Godlee *et al*, 2022). The interaction between these truncations and TMEM127 was assessed first. It was previously shown that mutations in transmembrane regions of SteD can prevent the interaction with TMEM127 (Alix *et al*, 2020). TMEM127 interacted with truncations of SteD comprising both transmembrane domains, together with either the N-terminal or C-terminal cytoplasmic domains. It did not interact with cytosolic SteD^1-41^ (the N-terminal 41 amino acid peptide) and there was little interaction with SteD^37-102^ (which lacks both the 36 cytoplasmic N-terminal amino acids and nine cytoplasmic C-terminal amino acids) (Fig 5B). Therefore, in addition to the transmembrane regions of SteD, the N- or C-terminal of SteD contribute to the SteD-TMEM127 interaction.

**Figure 5.**
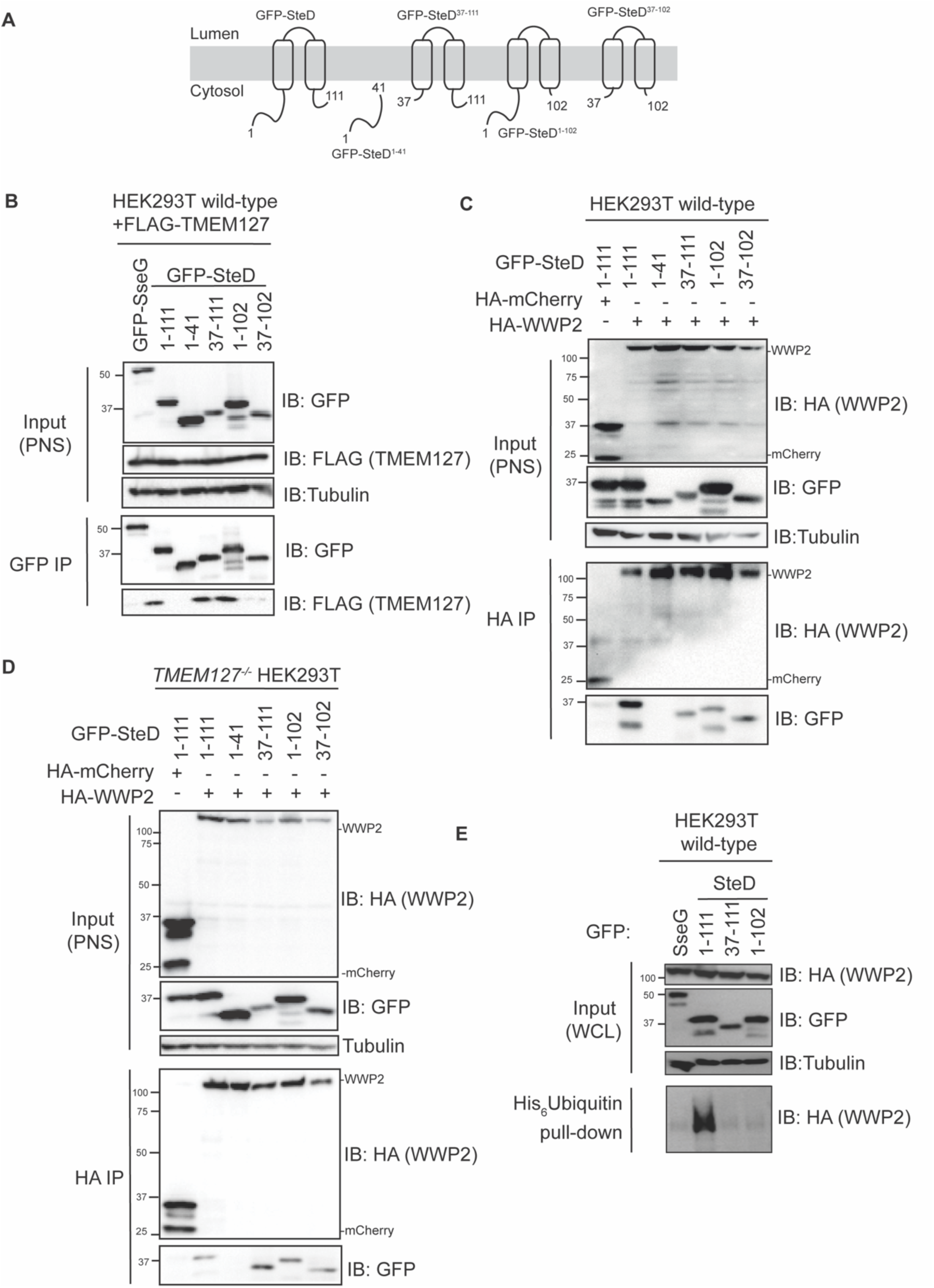
Truncations of SteD that interact with TMEM127 and WWP2 and disinhibit WWP2. A. A schematic of the GFP-SteD truncations used in these experiments. **B.** HEK239T cell transfection with GFP-SteD (WT or truncation) and FLAG-TMEM127, followed by GFP-immunoprecipitation (IP) and analysis of immunoprecipitate by immunoblotting. **C & D.** HEK239T cells (C. wild-type, D. TMEM127^-/-^) transfection with GFP-SteD (WT or truncation) and HA-WWP2, followed by HA-immunoprecipitation and analysis by immunoblotting. **E.** HEK293T cells transfection followed by His pulldown and analysis by immunoblotting. All blots are representative of at least three independent experiments. WCL, whole cell lysate. PNS, post-nuclear supernatant.

In the presence of TMEM127, SteD^37-111^, SteD^1-102^ and SteD^37-102^ interacted with WWP2 (Fig 5C). Since SteD^37-111^ and SteD^1-102^ interactions could be indirect (via TMEM127), we assessed SteD-WWP2 interactions in the absence of TMEM127. In this case, SteD^1-111^ (full length), SteD^37-111^, SteD^1-102^ and SteD^37-102^ all interacted with WWP2 (Fig 5D), providing strong evidence for an interaction between SteD and WWP2 that does not involve TMEM127. Despite being able to bind both TMEM127 and WWP2, SteD^37-111^ and SteD^1-102^ truncations were poor disinhibitors of WWP2 (Fig 5E). This suggests that the N- and C-terminal cytosolic regions of SteD are important for WWP2 disinhibition.

### SteD interacts with the C2 domain of WWP2 and the C2 domain contributes to disinhibition of WWP2

Since SteD does not have any PY motifs, we hypothesised that it might interact with a domain other than the WW domains and that these interactions might account for how SteD disinhibits WWP2. Immunoprecipitation of HA-WWP2 truncations (Fig 6A), showed that SteD interacted with all truncations that contain the C2 domain of WWP2 and the C2 domain (amino acids 1-171) was sufficient for this interaction (Fig 6A). There was also a very weak interaction between SteD and WWP2 truncations containing the HECT domain (Fig 6A). Given that SteD interacted with WWP2 domains that TMEM127 did not interact with (Fig 2C), these C2 domain-SteD interactions are likely to be direct.

**Figure 6.**
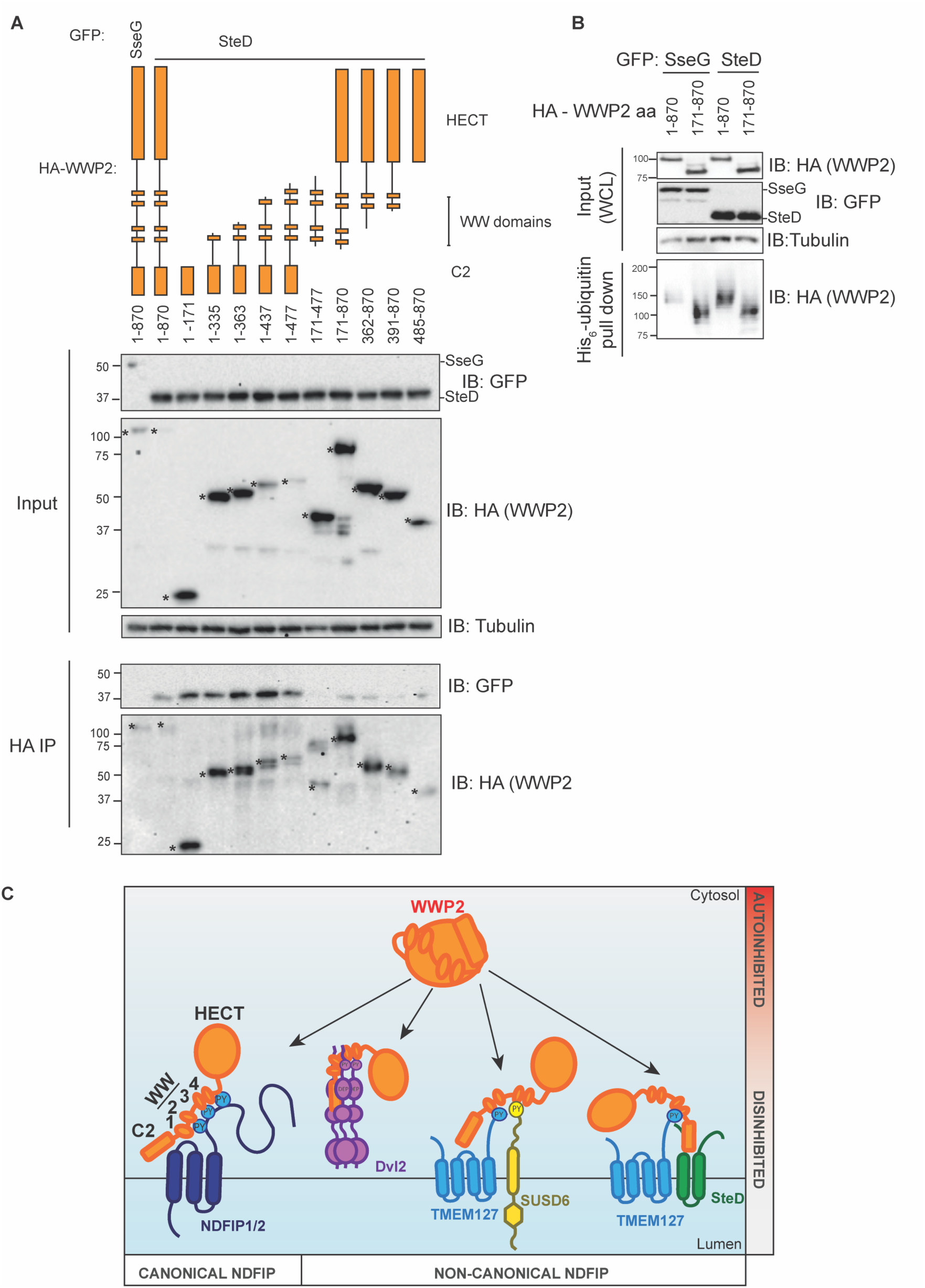
SteD interacts with the C2 domain of WWP2 and the C2 domain contributes to disinhibition of WWP2. A. HEK293T cell transfection with GFP-SseG or GFP-SteD and HA-WWP2 (full length or truncation), followed by HA-immunoprecipitation (IP) and analysis by immunoblotting. * Indicates relevant bands. **B.** HEK293T cell transfection followed by His pulldown and analysis by immunoblotting. **C.** Model of disinhibition of WWP2 by canonical and non-canonical Ndfips. All blots are representative of three independent experiments

The contribution of the C2 domain of WWP2 to its auto-inhibition is not clear. Wiesner et al, showed that *in vitro*, a WWP2 mutant lacking the C2 domain had a greater level of auto-ubiquitination compared to the wild-type protein (Wiesner *et al*, 2007). On the other hand, Chen et al showed that truncation of the C2 domain did not lead to increased auto-ubiquitination of WWP2 *in vitro* (Chen *et al*, 2017). We assessed the importance of the C2 domain to auto-inhibition of WWP2 with the His6-ubiquitin pull down assay in HEK293T cells by comparing full-length WWP2 with WWP2 lacking the C2 domain (WWP2^171-870^). In contrast to full-length WWP2, this truncation underwent disinhibition in the presence of the control *Salmonella* effector SseG, to a level similar to that detected when SteD was expressed together with either full length or C2-truncated WWP2 (Fig 6B). Our results are therefore in agreement with Wiesner et al (2017) and strongly suggest that binding of SteD to the C2 domain of WWP2 relieves its autoinhibition (Fig 6C).

## Discussion

Previous studies have shown that the NEDD4-like E3 ubiquitin ligases bind to PY motifs of Ndfips and this can lead to disinhibition of ligase activity. In this work we expand our understanding of requirements for adaptor binding and disinhibition of WWP2 by three mammalian proteins (TMEM127, NDFIP2 and SUSD6) and one from the pathogenic bacterium *Salmonella enterica* (SteD). Our experiments indicate that TMEM127, NDFIP2 and SUSD6 all interact with the WW2 domain of WWP2. SUSD6 also interacts with the WW3 and WW4 domains, and SteD only binds to the C2 domain. SteD and SUSD6 bind to WWP2 in the absence of TMEM127 but both SteD and SUSD6 require TMEM127 to disinhibit WWP2. Therefore, SUSD6/TMEM127 and SteD/TMEM127 function as co-disinhibitor pairs, which makes them a novel class of Ndfips. Intramembrane interactions between TMEM127 and SteD would enable formation of a three-protein complex and cooperative binding to both WW2 and C2 domains, leading to relief of autoinhibition and WWP2 disinhibition. It is not known if or how SUSD6 interacts with TMEM127, but together, their combined ability to bind multiple WW domains could enable WWP2 disinhibition. As TMEM127 is a flexible Ndfip able to function with both SUSD6 or SteD, there might be other endogenous TMEM127-partnering co-disinhibitors.

One PY motif of TMEM127 is required for stable binding to WWP2 (Alix *et al*, 2020), but we show here that it is not sufficient for WWP2 disinhibition. The three PY motifs of NDFIP2 are clustered within a region spanning amino acids 150 -186. Mund and Pelham found that a 52 amino acid peptide of NDFIP2 including the three PY motifs was able to disinhibit the related NEDD4 ligase Itch. All three PY elements were shown to contribute to activity (Mund & Pelham, 2009), and it was concluded that higher avidity of multiple PY–WW interactions is important for this. Our results on WWP2 disinhibition with PY point mutants of NDFIP2 follow the same pattern, but while the single PY3 and double PY1/PY2 point mutants had reduced binding to WWP2, the binding of single PY1 and PY2 point mutants to WWP2 was similar to that of the wild-type protein. Furthermore, both single PY1 and PY2 point mutants were markedly reduced in their ability to disinhibit WWP2, suggesting that PY1 and PY2 enable catalytic disinhibition, in addition to their partially redundant roles in binding WWP2. Another intriguing result from experiments with NDFIP2 was that despite its three PY motifs, full length NDFIP2 only bound to truncations of WWP2 containing the WW2 domain. Perhaps an initial strong interaction between WW2 and PY3 facilitates subsequent weaker (and undetectable by methods used here) interactions between PY1/PY2 and other WW domains. Structural and other biophysical/molecular dynamic studies are now required to show more precisely how these WW-PY interactions occur. These may also help to explain how TMEM127 and NDFIP2 interact with the same WW2 domain yet differ with respect to disinhibition of WWP2.

That TMEM127, SUSD6 and NDFIP2 interact with the WW2 domain of WWP2 suggests an important function for this domain. Structural work has shown that WW2 overlies a region called the hinge-loop region, which provides a point of flexibility between the N- and C-lobes of the HECT domain and is required for the ubiquitin transferase activity of the HECT domain (Chen *et al*, 2017; Verdecia *et al*, 2003). Therefore, binding of an Ndfip to the WW2 domain might contribute to disinhibition by preventing interactions between WW2 and this region of the HECT domain.

Previous work has shown that WWP1, WWP2 and Itch are autoinhibited by interactions between the HECT domain and the WW domains or regions between the WW domains (Chen et al., 2017; Jiang et al., 2019). In contrast, other NEDD4-like E3 ubiquitin ligases, such as NEDD4 and Smurf, are held in an autoinhibitory state through C2-HECT domain interactions (Weisner et al., 2007; Wang et al., 2010). It has been shown recently that C2 domain-mediated autoinhibition of Nedd4L ligase can be relieved by membrane curvature induced by a Bin-Amphiphysin-Rsv (BAR) domain protein, FCHO2, through an interaction between the C2 domain and the membrane (Sakamoto *et al*, 2024). In contrast to previous work assessing the contribution of the C2 domain to WWP2 autoinhibition *in vitro* (Chen et al., 2017; Wiesner et al., 2007), our assay assessed WWP2 disinhibition in the presence of known and unknown endogenous Ndfips and other factors, such as membrane association, that are known to contribute to WWP2 disinhibition. These factors might account for the differences between our results and those of Chen et al (2017). Our work shows that the C2 domain is required for autoinhibition of WWP2 and through its interaction with SteD, the autoinhibitory effect of the C2 domain is lost. This strongly suggests that the interaction between membrane-bound SteD and the C2 domain is allosteric or initiates other downstream events, such as protein aggregation, that releases the ligase activity of WWP2. Therefore, there appear to be at least two inhibitory interactions, involving both WW and C2 domains, that contribute to WWP2 autoinhibition.

Disinhibition of WWP2 by TMEM127 and SteD has similarities to the eukaryotic protein Dvl2. Dvl2 has a single PPxY motif that contributes to Dvl2-WWP2 binding but Dvl2 also interacts with the C2 domain of WWP2 via its Dishevelled, Egl-10 and Pleckstrin (DEP) domain. Furthermore, the DEP domain also contributes to the disinhibition of WWP2 (Mund *et al*, 2015). Although there are no sequence similarities between the DEP domain of Dvl2 and SteD, it is striking that Dvl2 interacts with WWP2 through a PPxY motif together with a C2-interacting/activating domain. Therefore, SteD and TMEM127 might be functionally analogous (as a pair) to Dvl2.

NDFIP1 and NDFIP2 were the first adaptors shown to disinhibit WWP2 through an interaction via their three PY motifs (Mund & Pelham, 2009). These could be considered canonical Ndfips. However, our findings along with previous work (Mund *et al*, 2015) show that Ndfips are more diverse in the way in which they interact with and disinhibit WWP2. The non-canonical Ndfips include (a) the cytoplasmic Dvl2 with its single PPxY motif, requirement for polymerisation, and interaction with the C2 domain (Mund *et al*, 2015); and those described here: (b) the SUSD6/TMEM127 endogenous co-activating pair, both containing PY motifs; and (c) the SteD/TMEM127 co-activating pair with a single PPxY motif of TMEM127 and C2-interacting properties of SteD (Fig 6C). Further structural work is now required to provide deeper mechanistic insights into the binding and disinhibition of WWP2 by these proteins.

## Acknowledgements

We thank Teresa Thurston for helpful comments on the manuscript and gift of the PX330 plasmid. We thank Camilla Godlee and Ondrej Cerny for advice and discussions. We are grateful to Hugh Pelham who provided the His_6_-ubiquitin plasmid.

## Author contributions

Samkeliso V. Blundell - Conceptualization, Investigation, Supervision, Writing Mei Liu – Investigation Romina Tocci - Investigation Andrea Majstorovic - Investigation Solène Tsang – Investigation David W. Holden - Conceptualization, Funding acquisition, Supervision, Writing

---

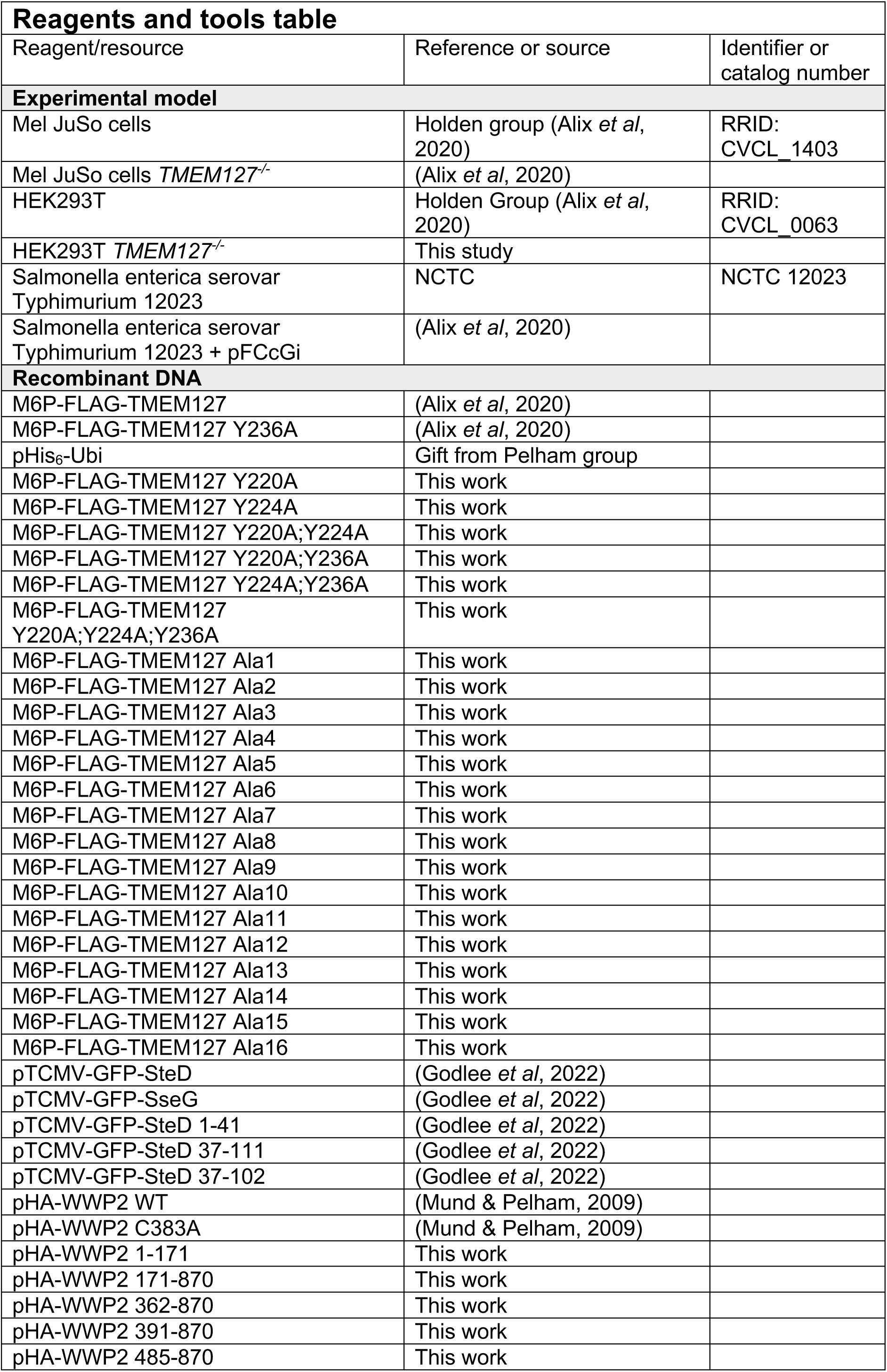
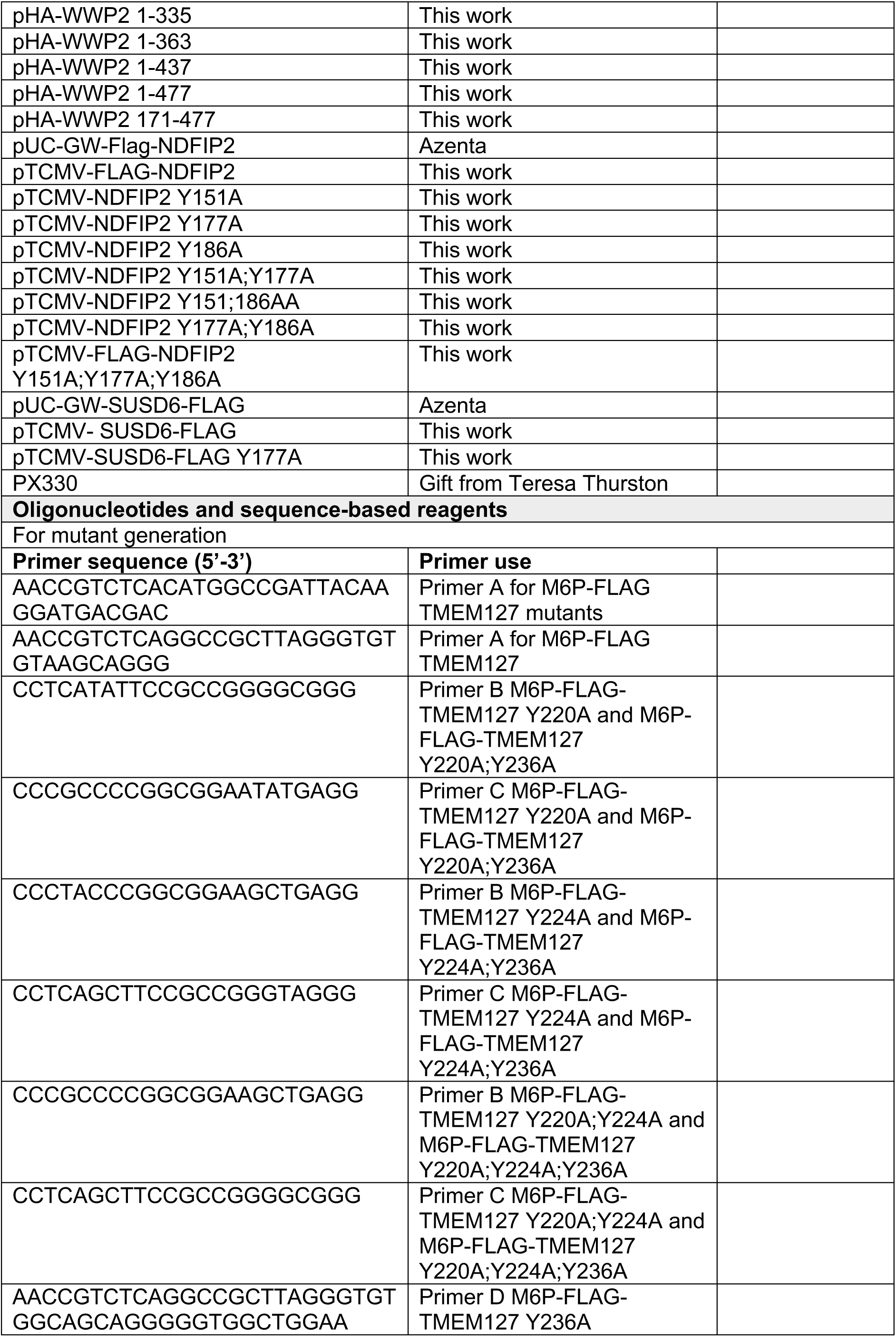
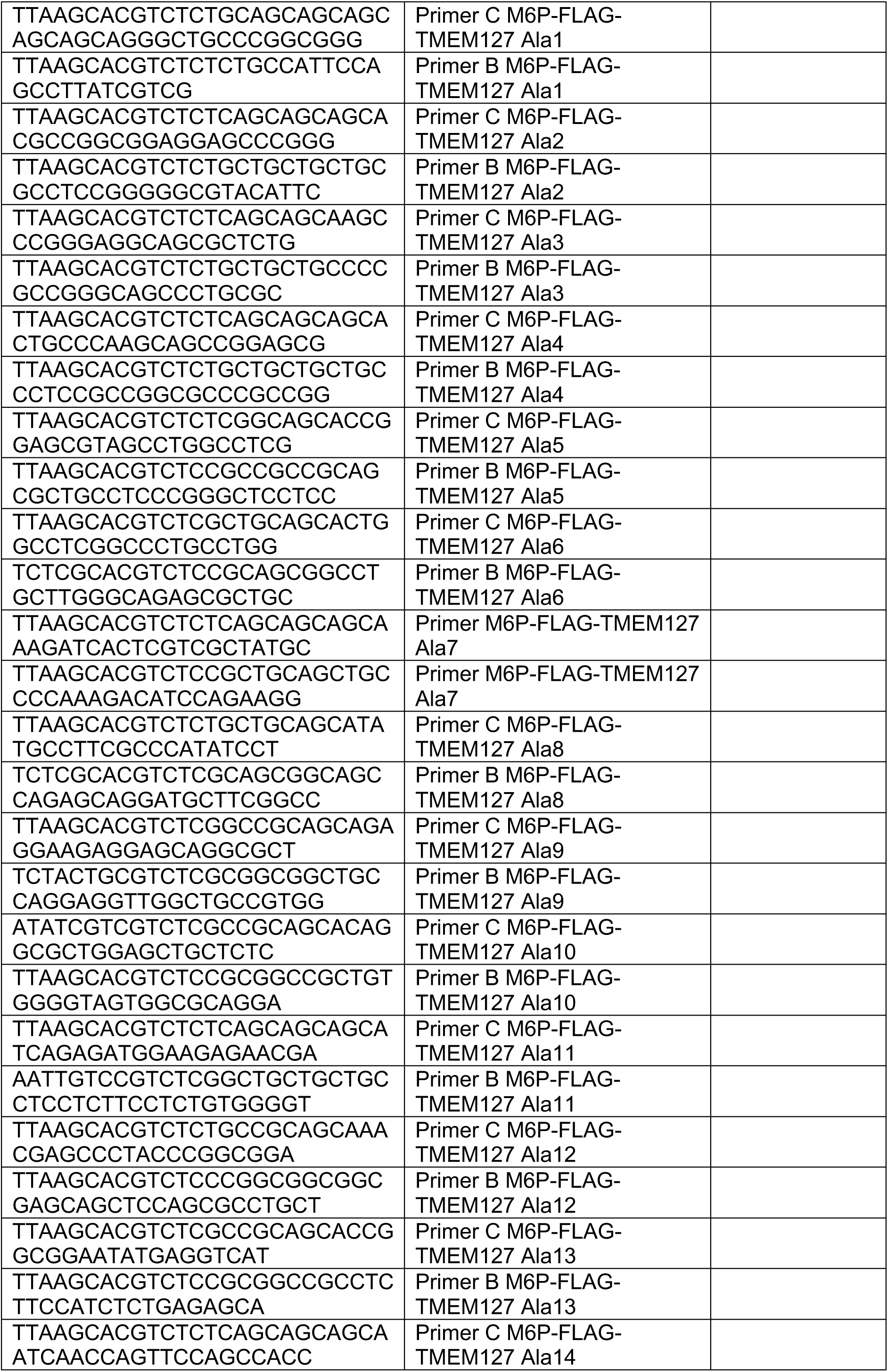
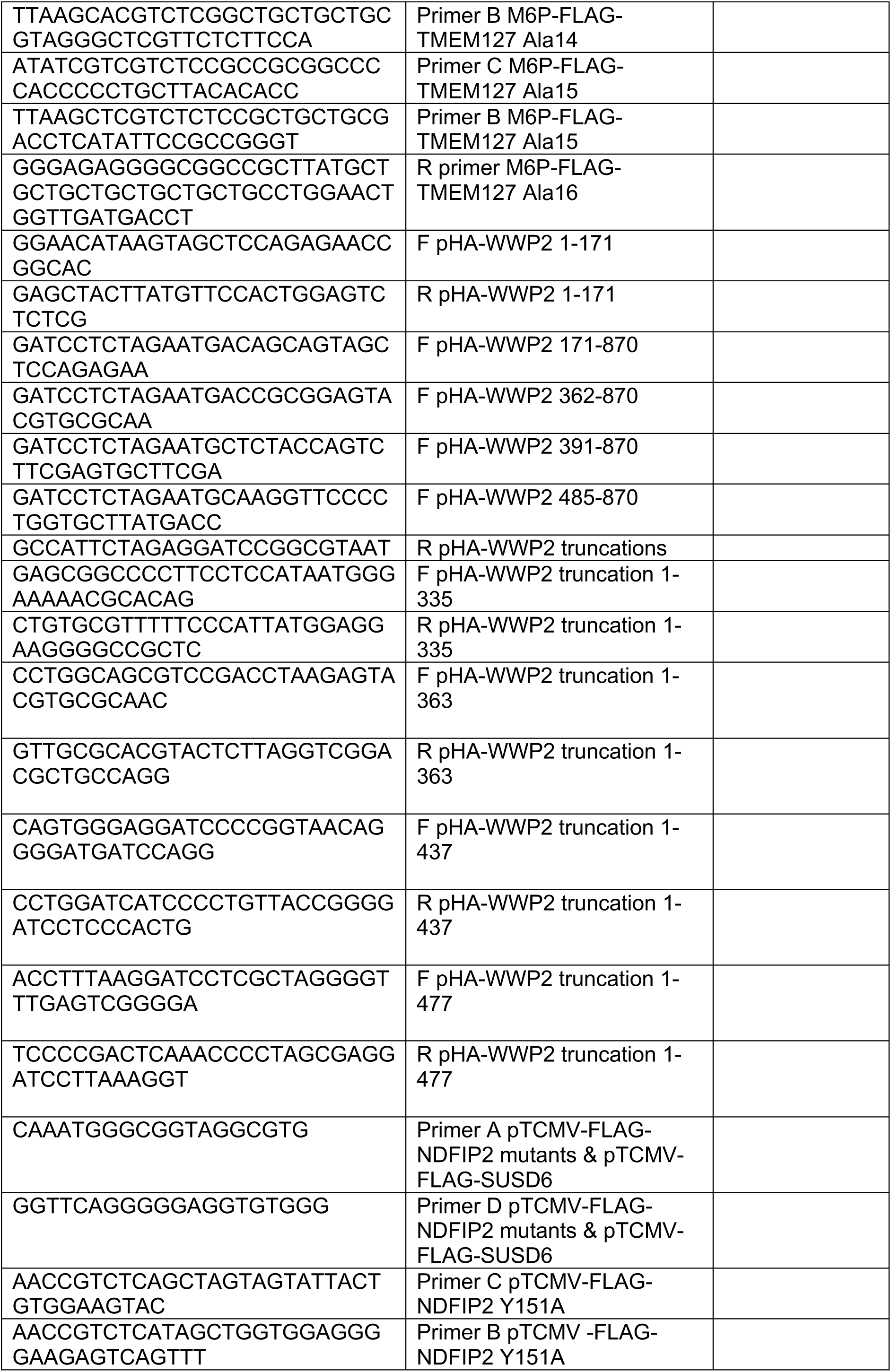
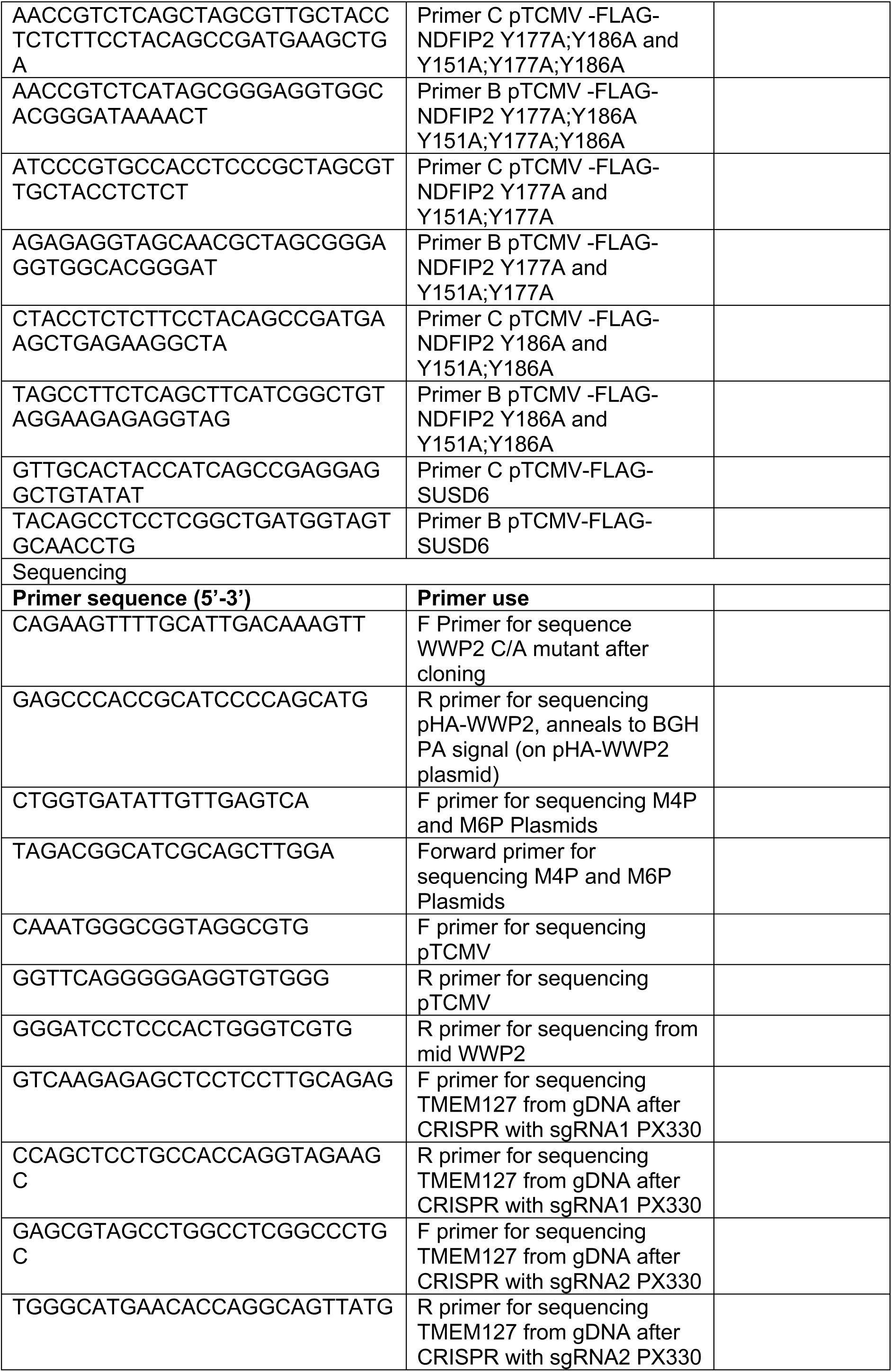
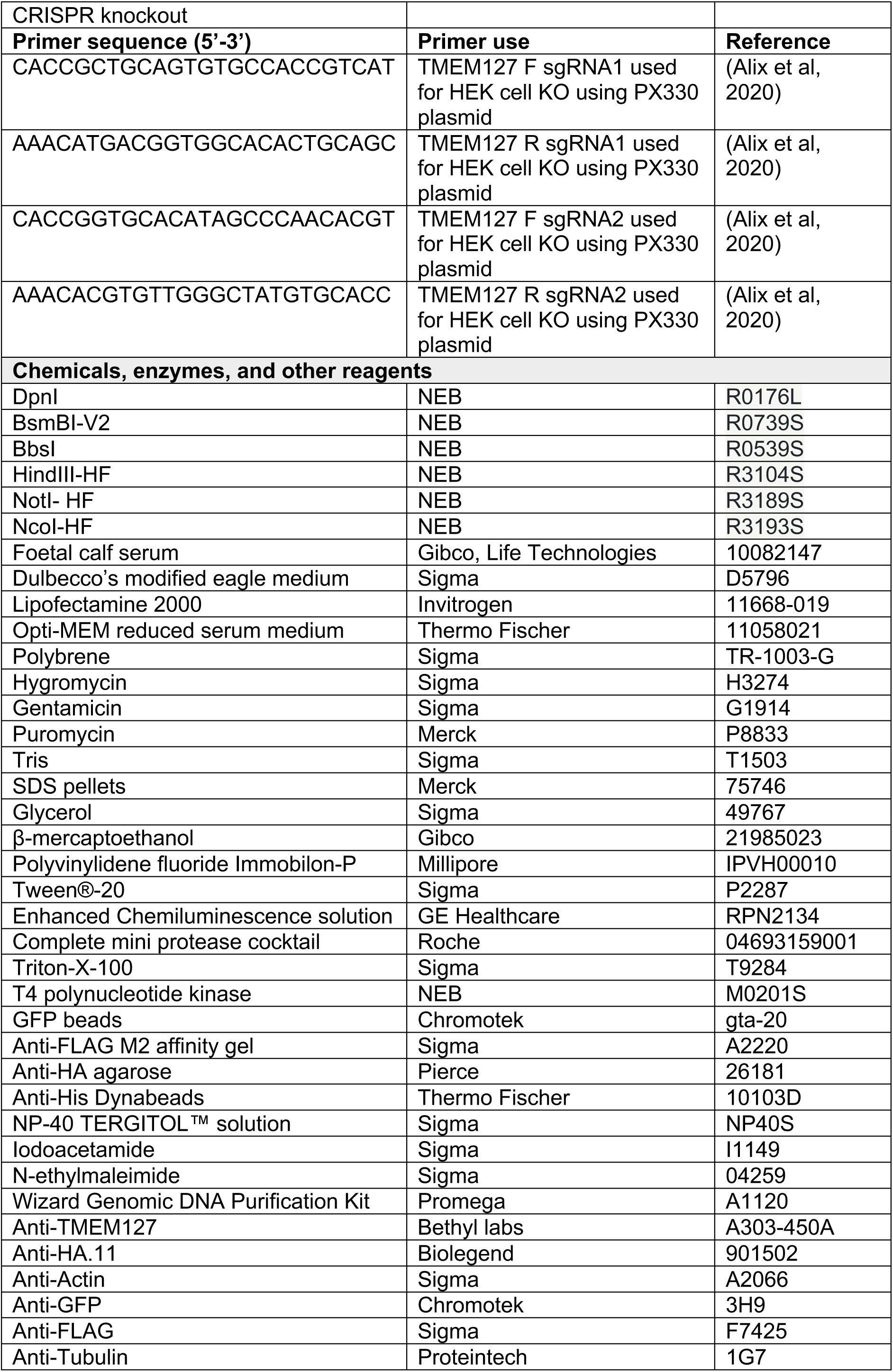
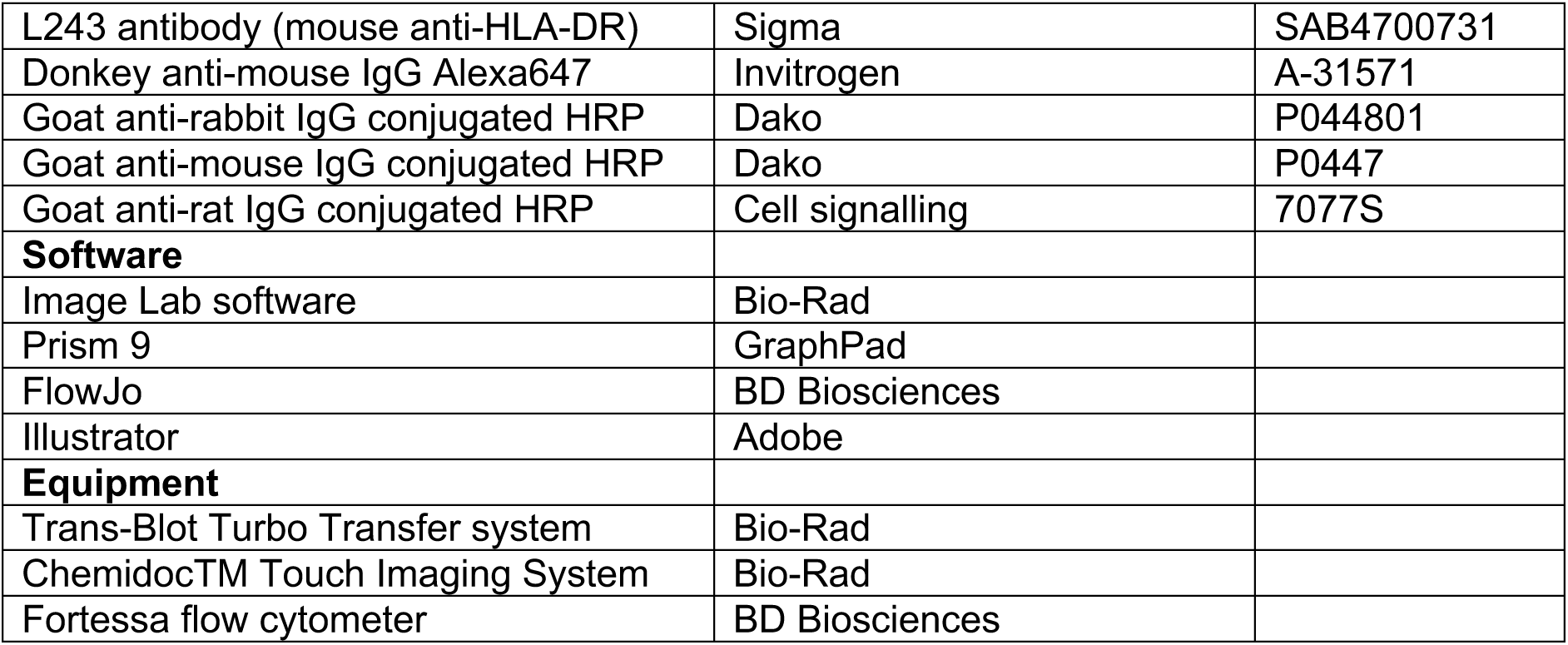

## Methods

### Bacterial strains

*Salmonella enterica* serovar Typhimurium 12023 was used for infections. *Escherichia coli* strain DH5α was used for transformation of plasmids to produce bacterial stocks. Bacteria were grown in Luria-Bertani (LB) medium supplemented with carbenicillin (50 μg ml^-1^), kanamycin (50 μg ml^-1^) or chloramphenicol (30 μg ml^-1^) as appropriate.

### Plasmid construction

Mutations were created using the overlap-PCR method (Heckman & Pease, 2007) or by one-step site-direct mutagenesis(Liu & Naismith, 2008). Following one-step site-directed mutagenesis, the PCR product was then restricted with DpnI. The ligated or DpnI restricted products were transformed into *E. coli* DH5α. Colonies were selected for colony PCR and then sent for sequencing.

### Cell culture

Human Embryonic Kidney 293T (HEK293T) cells and human Mel JuSo cells were maintained in Dulbecco’s modified eagle medium (DMEM) supplemented with 10% heat-inactivated foetal calf serum (FCS) cells were incubated at 37 °C in 5% CO2. When new cell lines were being created, 50 U ml^-1^ of penicillin and 50 μg ml^-1^ streptomycin was added to the DMEM.

### Transfection and virus production

For transfections, HEK293T or Mel JuSo cells were seeded. 16 h later, cells were transfected with a total of 0.25 – 1 μg DNA plus 3-5 μl of lipofectamine 2000 in Opti-MEM reduced serum medium per well. Cells were then collected 16-20 h later. For virus production, HEK293T cells were co-transfected with lentiviral expression vector and packing plasmids VSVG and GagPol. The lentiviral expression vectors used were M6P encoding FLAG-TMEM127 (PY mutants and alanine mutants). The culture media was replaced 24 h post-transfection and supernatant containing viruses was collected at 48 h post-transfection and filtered.

### Generation of stable cells lines

To generate Mel JuSo cells stably expression FLAG-TMEM127 for alanine scanning mutagenesis, lentiviruses were added to Mel JuSo cells with polybrene (8 μg ml-1). At 48 h post-transduction cells were selected for with hygromycin (800 μg ml-1).

### Infection

*S*. Typhimurium strains were incubated overnight at 37 °C in a shaking incubator with the appropriate antibiotics. The overnight culture was diluted 1:33 in LB and incubated in a shaking incubator at 37 °C for 3.5 h before adding to Mel JuSo cells at an MOI of 1:100 for 30 min. Cells were washed twice and incubated with fresh DMEM containing gentamicin (100 μg ml^-1^) for 1 h. After 1 h, the media was exchanged with fresh media containing gentamicin (20 μg ml^-1^). Cells were processed at 20 h post-invasion.

### Cell lysis for western immunoblotting

Cells were washed in PBS and incubated in trypsin for 1 min at 37 °C. Cells were washed twice in PBS and then lysed in 0.1% triton in PBS for 10 min on ice before centrifugation at 13,000 revolutions per minute (rpm) at 4 °C for 10 min. The post-nuclear supernatants (PNS) were collected and boiled in 4x SDS loading buffer (0.25 mM Tris pH 6.8, 10% SDS, 50% glycerol, 5% β-mercaptoethanol) at 95 °C for 5 min.

### SDS-PAGE and western immunoblotting

Cell lysates in SDS buffer were loaded and run on 8-12% polyacrylamide gels. Protein was transferred on to polyvinylidene fluoride (PVDF) membranes by semi-dry transfer using the Trans-Blot Turbo Transfer system. Membranes were blocked with 5% milk in Tris buffered saline with 0.01% Tween® (TBS-T) for 1 h at room temperature. Membranes were incubated overnight with the primary antibody in 5% milk in TBS-T at 4°C. The following morning membranes were three times in TBS-T for a total of 30 min, incubated with secondary antibody in 5% milk in TBS-T for 90 min and washed three times in TBS-T for a total of 30 min. Enhanced Chemiluminescence solution was then applied and detection of signal using a ChemidocTM Touch Imaging System.

### Green fluorescent protein (GFP), FLAG and haemagglutinin (HA) co-immunoprecipitation (IP)

HEK293T cells transfected with plasmids outlined in Figures or stable cell lines were collected and the IP lysis solution (0.5% Triton-X-100, 150 mM NaCl, 50 mM Tris-Cl pH 7.4, 5 mM EDTA, 5% glycerol, complete mini protease cocktail) was added. Samples were lysed and then centrifuged at 16,000 g for 10 min at 4°C. Post-nuclear supernatant (PNS; input) was mixed with of washed GFP beads, anti-FLAG gel or anti-HA agarose for 2 h at 4°C on a 360° roller. Beads/agarose were then washed 3-5 times in wash buffer (0.5% Triton-X-100, 50 mM NaCl, 50 mM Tris-Cl pH 7.4, 5 mM EDTA, 5% glycerol). For the GFP IP after the final wash, beads were resuspended in SDS loading buffer. For the HA and FLAG IP, the gel or agarose was resuspended in SDS loading buffer without the reducing agent. All samples (input and immunoprecipitate) were boiled at 95 °C for 5 minutes before analysis by SDS-PAGE and immunoblots carried out as outlined above.

### His pull down of His-tagged ubiquitin

HEK293T cells were co-transfected with plasmids encoding proteins of interest and a plasmid encoding His6-tagged ubiquitin (Treier *et al*, 1994; Mund & Pelham, 2009). 16 h later, cells were lysed directed in a denaturing lysis/wash buffer (8 M urea, 50 mM Tris pH 8, 2 mM N-ethylmaleimide, 10 mM IAA, 0.2% SDS, 0.5% NP40). The lysate was sonicated. 20 μl of Dynabeads per sample were washed twice in the lysis/wash buffer. The lysate was incubated with dynabeads on a rolling incubator for 2 h at room temperature. Dynabeads were then washed three times in the lysis/wash buffer and finally resuspended in SDS loading buffer. The input and beads were then boiled at 95 °C for 5 min before analysis by SDS page and western blotting.

### CRISPR knock out of *TMEM127* using the PX330 plasmid

The PX330 plasmid encodes both the small guide RNA (sgRNA) and the Cas9 nuclease. The sgRNAs targeting the *TMEM127* gene were the same sequence used in a previous publication (Alix et al., 2020). The PX330 construct was then transfected in HEK293T cells. The sgRNA oligos were annealed and phosphorylated in one reaction step. The annealed oligos and PX330 plasmid were then restricted using BbsI and the resultant products were incubated with T4 ligase. Ligated product was transformed into *E. coli* DH5α as described above. HEK293T cells were transfected with TMEM127 sgRNA PX330 plasmid and selected for with puromycin (0.8 μg ml^-1^). Single cells were placed into a 96 well using the dilution method and grown on to create clonal populations. An aliquot of the cells was lysed and proteins levels of TMEM127 were assessed by SDS-PAGE and western blotting. Suitable cell lines were then selected for genome sequencing. Genomic DNA was extracted using the Wizard Genomic DNA Purification Kit and samples sent for sequencing.

### Flow cytometry

Surface levels of mMHCII on Mel JuSo cells were measured following infection with *S*. Typhimurium 12023 expressing GFP. Mel JuSo cells were infected as outlined above. 20 h post-invasion, cells were detached from cell culture plates using 2mM EDTA in PBS. All antibodies were diluted in FACS buffer (5% FCS and 1mM EDTA in PBS). Cells were incubated with monoclonal L243 for human mMHCII for 30 min on ice, washed in cold PBS and then incubated with Alex Fluor 647 donkey anti-mouse for 30 min on ice. Cells were then resuspended in FACS buffer and taken for immediate analysis by flow cytometry. Data were acquired using a Fortessa flow cytometer; the fluorescence intensity of each fluorophore for each cell was measured. Data were analysed using FlowJo v10 software.

The geometric mean fluorescence intensity (gMFI) was then calculated. Surface levels of mMHCII were calculated as a gMFI of infected (GFP-positive) over gMFI of non-infected cells (GFP-negative).

### Data analysis: graphs and statistics

The graph was created and flow cytometry data analysed using GraphPad Prism 9. Multiple values were all compared against each for analysis of flow cytometry data using an Ordinary one-way ANOVA followed by Tukey’s multiple comparisons test.

