## Supplemental Figures 1-3 for "*Salmonella* SteD and mammalian SUSD6 use TMEM127 to co-disinhibit the WWP2 E3 ubiquitin ligase"

| | | | | | | | |
|---|---|---|---|---|---|---|---|
| **Wild type** | 140 | ALETDSSPPP | YSSITVEVPT | TSDTEVYGEF | YPVPPPYSVA | TSLPTYDEAE | 190 |
| **PY1(Y151A)** | 140 | ALETDSSPPP | ASSITVEVPT | TSDTEVYGEF | YPVPPPYSVA | TSLPTYDEAE | 190 |
| **PY2 (Y177A)** | 140 | ALETDSSPPP | YSSITVEVPT | TSDTEVYGEF | YPVPPPASVA | TSLPTYDEAE | 190 |
| **PY3 (Y186A)** | 140 | ALETDSSPPP | YSSITVEVPT | TSDTEVYGEF | YPVPPPYSVA | TSLPTADEAE | 190 |
| **PY12 (YY151,177AA)** | 140 | ALETDSSPPP | ASSITVEVPT | TSDTEVYGEF | YPVPPPASVA | TSLPTYDEAE | 190 |
| **PY13 (YY151,186AA)** | 140 | ALETDSSPPP | ASSITVEVPT | TSDTEVYGEF | YPVPPPYSVA | TSLPTADEAE | 190 |
| **PY23 (YY177,186AA)** | 140 | ALETDSSPPP | YSSITVEVPT | TSDTEVYGEF | YPVPPPASVA | TSLPTADEAE | 190 |
| **PY123(YYY151,177,186AAA)** | 140 | ALETDSSPPP | ASSITVEVPT | TSDTEVYGEF | YPVPPPASVA | TSLPTADEAE | 190 |

**S1 NDFIP2**

**A.** Amino acid sequence of NDFIP2 showing PY mutants highlighted in red.

A

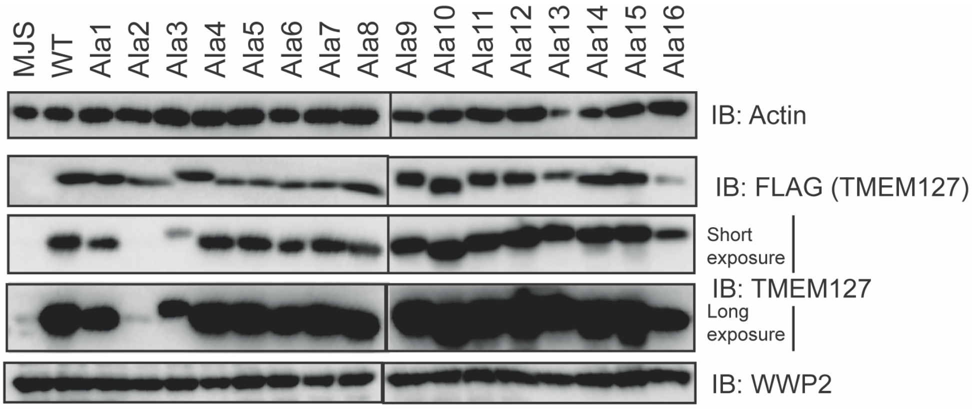


B

| | |
|---|---|
| **WT** | 210 LSEMEENEPY PAEYEVINQF QPPPAYTP 238 |
| **Y220A** | 210 LSEMEENEPA PAEYEVINQF QPPPAYTP 238 |
| **Y224A** | 210 LSEMEENEPY PAEAEVINQF QPPPAYTP 238 |
| **Y236A** | 210 LSEMEENEPY PAEYEVINQF QPPPAATP 238 |
| **Y220A;Y224A** | 210 LSEMEENEPA PAEAEVINQF QPPPAYTP 238 |
| **Y220A;Y236A** | 210 LSEMEENEPY PAEAEVINQF QPPPAATP 238 |
| **Y224A;Y236A** | 210 LSEMEENEPY PAEAEVINQF QPPPAATP 238 |
| **Y220A;Y224A;Y236A** | 210 LSEMEENEPA PAEAEVINQF QPPPAATP 238 |

C

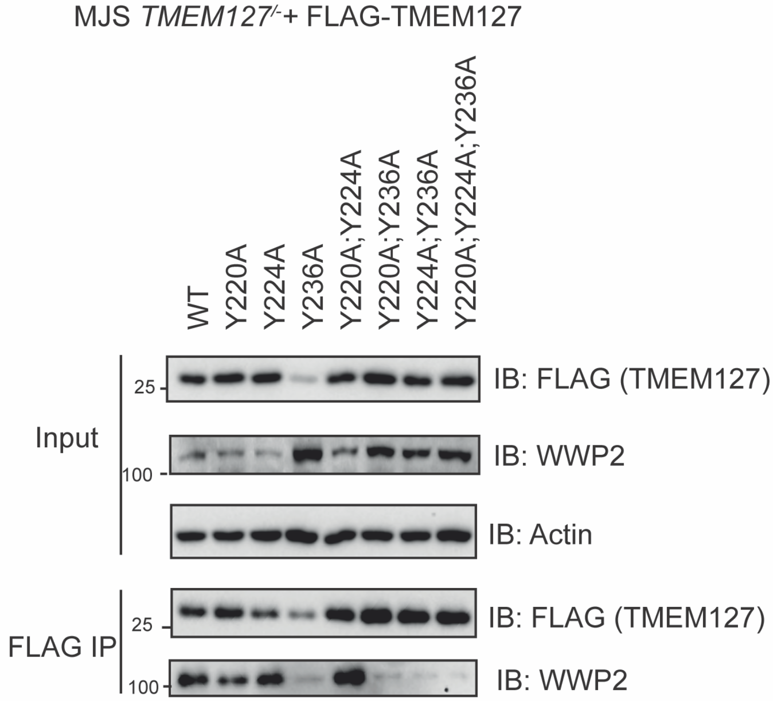


**S2 TMEM127**

**A**. Mel JuSo cells or mutant TMEM127 alanine mutant (as indicated) cell line levels of FLAG-TMEM127 as assessed by immunoblotting. **B.** Amino acid sequence of TMEM127 and PY mutants created from 210 to 238 with tyrosines within possible PY motifs indicated in red and mutations created indicated in orange. **C.** FLAG immunoprecipitation of TMEM127 PY mutant cell line (as indicated) and analysis of immunoprecipitate by immunoblotting. The blot shown is representative of the three independent experiments.

A

| SteD truncation | Amino acids | Subcellular localisation |
|---|---|---|
| Full length | 1-111 | Normal (Golgi and endosomes) |
| N-terminal cytoplasmic region | 1-41 | Cytoplasmic |
| N-terminal truncation | 37-111 | Plasma membrane, Golgi and endosomes |
| C-terminal truncation | 1-102 | Normal (Golgi and endosomes) |
| N- & C-terminal truncation | 37-102 | Plasma membrane, Golgi and endosomes |

B

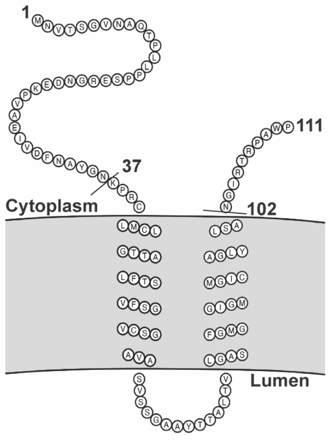


**S3 SteD**

**A.** A summary of GFP-SteD truncations used in Figure 5, truncations previous published in (Godlee *et al*, 2022). **B.** Schematic of truncations of SteD in relation to SteD topology
